# A monoclonal antibody collection for *C. difficile* typing

**DOI:** 10.1101/2023.09.21.557191

**Authors:** Lise Hunault, Patrick England, Frédéric Barbut, Bruno Iannascoli, Ophélie Godon, François Déjardin, Christophe Thomas, Bruno Dupuy, Chunguang Guo, Lynn Macdonald, Guy Gorochov, Delphine Sterlin, Pierre Bruhns

**Affiliations:** Institut Pasteur, Université Paris Cité, INSERM UMR1222, Antibodies in Therapy and Pathology, 75015 Paris, France; Sorbonne Université, INSERM, CNRS, Centre d’Immunologie et des Maladies Infectieuses (CIMI-Paris), 75013 Paris, France; Sorbonne Université, Collège doctoral, 75005 Paris, France; Institut Pasteur, Université Paris Cité, CNRS UMR3528, Plateforme de Biophysique Moléculaire, 75015 Paris, France; National Reference Laboratory for Clostridium difficile, 75012 Paris, France; Université Paris Cité, INSERM UMR-1139, Paris, France; Production and Purification of Recombinant Proteins Facility, Institut Pasteur, 75015 Paris, France; Institut Pasteur, Université Paris-Cité, UMR-CNRS 6047, Laboratoire Pathogenèse des Bactéries Anaérobies, 75015 Paris, France; Regneron Pharmaceuticals, Tarrytown, NY, USA

**Author notes:** shared senior authorship (Guy Gorochov, Delphine Sterlin, Pierre Bruhns).

**Keywords:** *Clostridioides difficile*, monoclonal antibodies, S-layer, hybridomas

## Abstract

*Clostridioides difficile* is the leading cause of antibiotic-associated diarrhea and pseudomembranous colitis in adults. Various *C. difficile* strains circulate currently, associated with different outcomes and antibiotic resistance profiles. However, most studies still focus on the reference strain 630 that does not circulate anymore, partly due to the lack of immunological tools to study current clinically important *C. difficile* PCR ribotypes. Herein, we immunized mice expressing human variable antibody genes with the Low Molecular Weight (LMW) subunit of the surface layer protein SlpA from various *C. difficile* strains. Monoclonal antibodies purified from hybridomas bound LMW with high-affinity and whole bacteria from current *C. difficile* ribotypes with different cross-specificities. This first collection of anti-*C. difficile* mAbs represent valuable tools for basic and clinical research.

## INTRODUCTION

*Clostridioides (*formerly *Clostridium) difficile* is an anaerobic, gram-positive, and spore-forming bacterium that is the main agent responsible for antibiotic-associated diarrhea and pseudomembranous colitis in adults ^1^. In the past decades, there was a drastic increase in the incidence of both healthcare-associated *C. difficile* infection (CDI) and community-acquired CDI^2^. There is a large phylogenetic diversity of *C. difficile* with more than 300 distinct PCR-ribotypes (RT) reported worldwide, including hypervirulent lineages associated with increased transmission and mortality^3–6^. The latest epidemiology data worldwide reported that 5 ribotypes i.e., RT001, RT002, RT014, RT027 and RT078, account for approximately 50% of the infections^7^.

Whereas several advances such as fluorescent mutants and novel fingerprinting techniques have contributed to a better understanding of *C. difficile* diversity and physiology^8– 10^, basic research still relies on one single strain *i*.*e*., *C. difficile* 630 that belong to RT012. An increasing number of studies has been performed on the hypervirulent ribotype 027, which caused major outbreaks in the United States and Europe at the end of the 2010s^11,12^. Other ribotypes remain largely unexplored even though some are associated with antibiotic resistance and increased severity^3^, which can be partly explained by the lack of genetic and immunological tools to study these strains.

*C. difficile* surface is composed of adhesins e.g., the flagellar cap protein FliD, the flagellin FliC, the cell wall protein Cwp66, the surface layer protein SlpA, and the protease Cwp84^13^. SlpA is expressed on the bacterial surface of all ribotypes and plays a crucial role in the pathogenesis and virulence of *C. difficile* by mediating interactions with the host cells and the surrounding environment^14–17^. SlpA contains two biologically distinct entities, the high-molecular weight (HMW) and the low molecular weight (LMW) subunits that assemble on the bacterial surface into a paracrystalline lattice^18^. Sequence variations of SlpA have been reported for the LMW that correlate with the diversity of clinical isolates, whereas the HMW is less variable^19,20^. SlpA is highly immunogenic, meaning it can trigger an immune response in the host^21^. Indeed, antibodies against SlpA have been detected in the sera of patients infected with *C. difficile*, indicating its potential as a target for vaccine development^21,22^.

In this work, we generated the first collection of mAbs that bind and discriminate predominant clinical ribotypes of *C. difficile*. Knock-in mice expressing human antibody variable genes for the heavy (V_H_) and light chain (V_L_)^23,24^ were immunized with a collection of recombinantly expressed LMW from five clinically relevant *C. difficile* ribotypes i.e., RT001, RT002, RT014, RT027 and RT078. Hybridomas were generated and their corresponding IgG mAbs bound both recombinant LMW *in vitro* and LMW naturally expressed on the bacterial surface. At least one mAb was identified against each of the five ribotypes used for immunization, with 6 mAbs being cross-reactive between LMW subunits of two different *C. difficile* ribotypes. The reduced sequence identity of LMW between different *C. difficile* ribotypes^25^ allows for specific identification of bacterial ribotypes by this anti-LMW mAb collection that represents a novel toolkit for *C. difficile* research.

## RESULTS

LMW SlpA subunits from 5 predominant ribotypes of *C. difficile* i.e., RT001, RT002, RT014, RT078 and RT027 (Fig. 1a), were recombinantly produced from transformed *Escherichia coli* as his-tagged soluble proteins and affinity-purified. As anti-LMW antibodies may potentially be of therapeutic interest for the treatment of CDIs, we used knock-in mice in which the endogenous genes encoding the heavy chain variable domain (VH) and the kappa light chain variable domain (Vκ) were replaced by their human counterparts (Velocimmune)^23,24^ with one modification, i.e., only one allele of the endogenous Vκ locus was replaced by human Vκ segments, and the second allele of the endogenous Vκ locus was replaced by human Vλ segments (Fig. 1b). As the Vk locus expresses 95% of the light chains in mice ^26^, placing human Vλ segments at the Vk locus increases the variability of light chain expression. Thus, after hybridoma identification, cloning of these VH and VL into vectors containing human heavy and light chain constant domains, allows for direct development - *in fine* – of fully human anti-LMW mAbs. To generate hybridomas, mice were immunized at D0, D21 and D42 with 50 μg/mouse of each LMW (Fig. 1c). High anti-LMW IgG serum titers were obtained in all mice at day 42 (Fig. 1d). Mice were boosted with all five LMW at equimolar ratio (Fig. 1c), and their spleen harvested 4 days later. Two different protocols were tested and gave similar results; one based on the similarity between the LMW – grouping two highly similar LMW in a single immunization; one based on their frequency in current CDI – grouping LMW corresponding to current clinical ribotypes in a single immunization (Supp. Fig. 1). More than 700 hybridomas were generated and among them 100 hybridoma were found to secrete anti-LMW antibodies.

**Figure 1:**
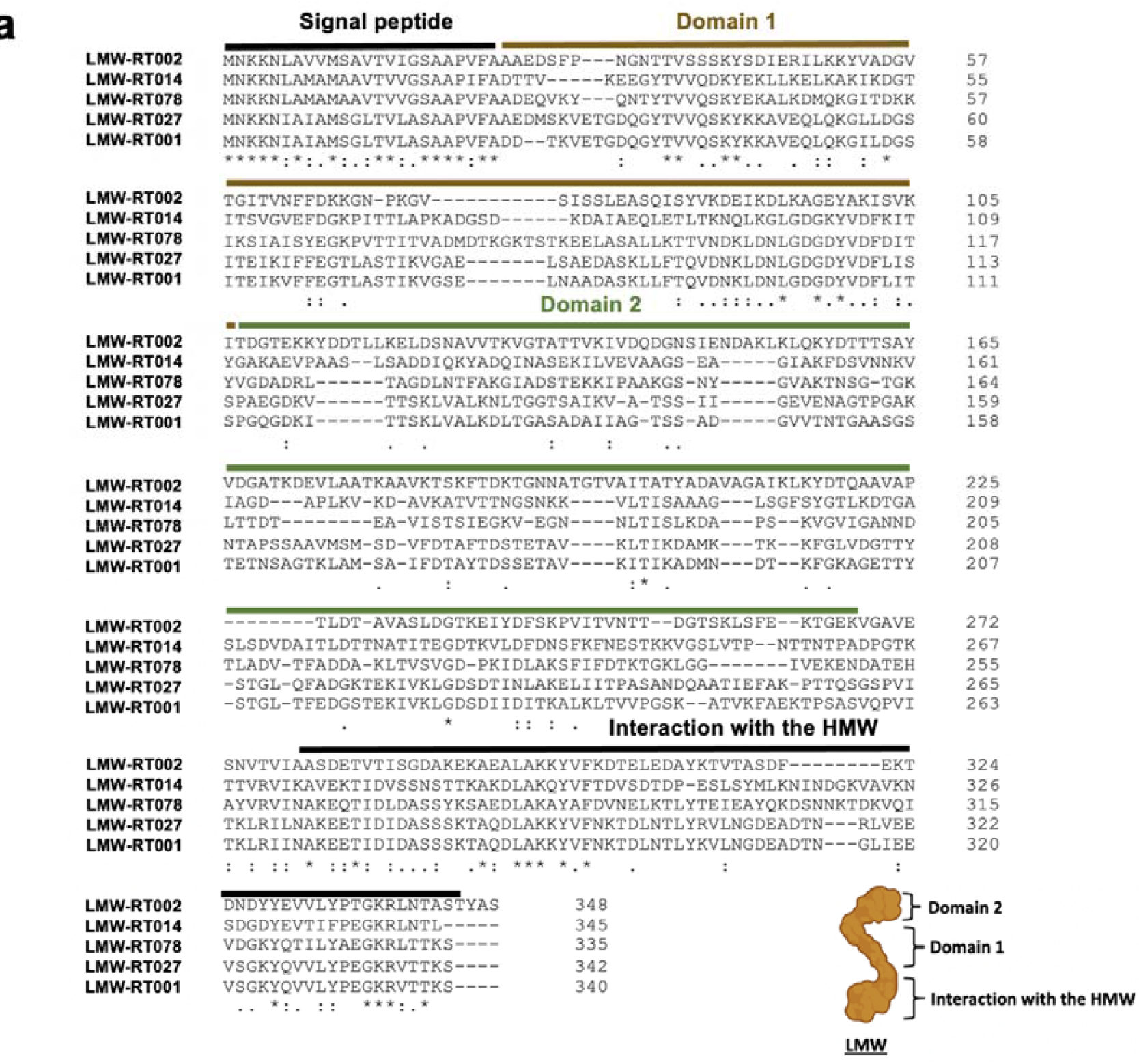
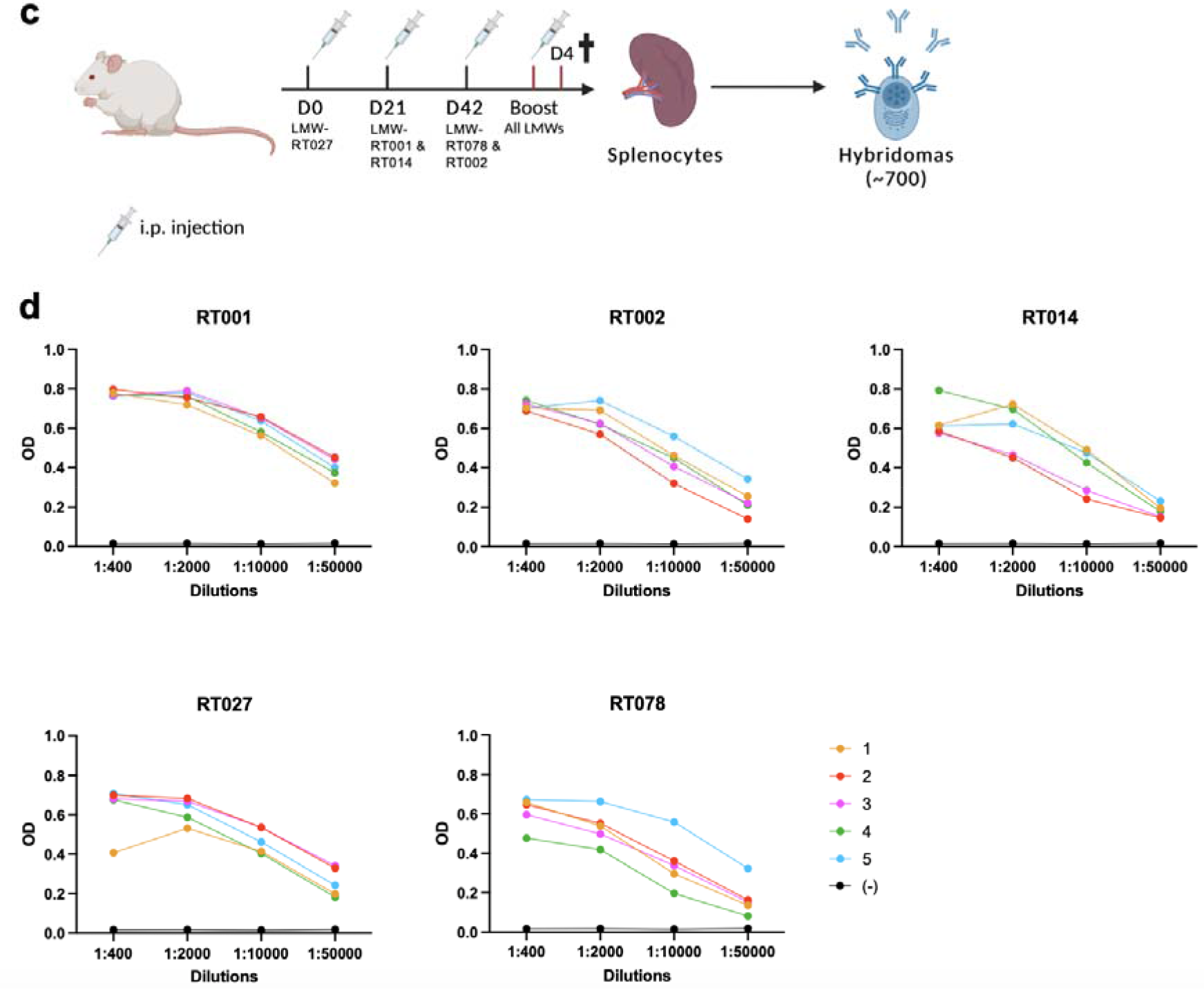
Generation of anti-LMW-specific hybridomas from immunized mice. (**a**) Sequence alignments of the LMW of five clinical ribotypes (LMW-RT001, LMW-RT002, LMW-RT014, LMW-RT027, LMW-RT078) by ClustalOmega software. Fully conserved residues are indicated by ^*^, groups of strongly similar properties by ^:^ and groups of weakly similar properties by *,. or :. Signal peptide, domain 1 and 2 and the domain that interacts with the HMW are indicated. (**b**) Schematic of the generation of mice knock-in for the human variable VDJ segments in the endogenous variable heavy chain locus, and for the human variable VJ segments in the endogenous variable light chain kappa locus. (**c**) Protocol outline. Mice were immunized with LMW proteins according to the represented scheme combined to alum and *Bordetella pertussis* toxin. Four days after the last boost, spleens were collected and hybridoma generated. (**d**) Sera titers at day 42 of immunized mice for recombinant LMW-RT001, LMW-RT002, LMW-RT014, LMW-RT078, LMW-RT027 measured by ELISA. OD values for several dilutions for mice #1 to #5 are represented. Black curves (-) represent sera titers of a naive mouse.

Among these 100 hybridomas, the 14 clones displaying the highest ratio of LMW binding by ELISA compared to IgG concentration in their culture supernatant were expanded and their antibodies purified. Their binding profiles towards the five recombinant LMW proteins were assessed by ELISA (Fig. 2a). 12 out of 14 (86%) significantly bound LMW-RT001 with variable profiles, 1 out of 14 (7%) bound LMW-RT002, 1 out of 14 (7%) bound LMW-RT014, 6 out of 14 (43%) bound LMW-RT078 and 11 out of 14 (78%) bound LMW-RT027. Among the eleven LMW-RT027-binding mAbs, four (36%) cross-reacted strongly with LMW-RT001 (mAb SG8, TF1, TH4 and VA10) and one with both LMW-RT001 and LMW-RT078 (mAb RF12). mAb QE2 cross-reacted with four LMWs: LMW-RT001, LMW-RT014, LMW-RT027 and LMW-RT078. Among the three mAbs that did not recognize LMW-R0T27, mAb RA11 was specific for LMW-R0T78, mAb UA5 cross-reacted with LMW-RT001 and LMW-RT002, and mAb SC6 cross-reacted with LMW-RT001 and LMW-RT078 (Fig. 2b).

**Figure 2:**
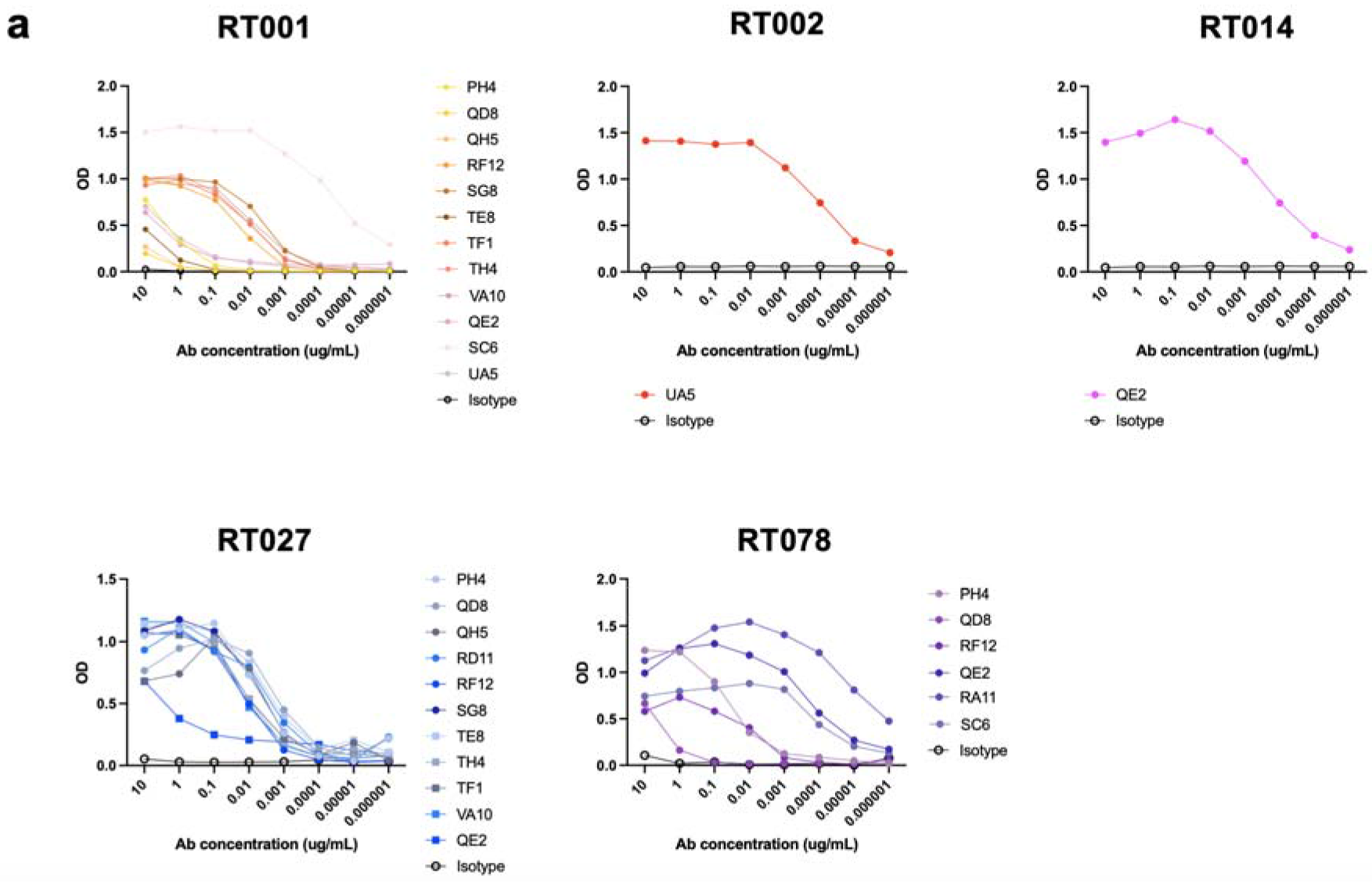
Specificities of anti-LMW mAbs. ELISA results (OD values 492 nm versus 620 nm) against recombinant LMW-RT001, LMW-RT002, LMW-RT014, LMW-RT078 and LMW-RT027 of IgG mAbs at indicated concentrations. Black curves represent isotype controls.

We next evaluated the affinity of the mAbs displaying the strongest interactions with their respective targets i.e., LMW-RT001, LMW-RT002, LMW-RT014, LMW-RT078 and LMW-RT027, by Bio Layer Interferometry (BLI), coupling IgGs to the sensors and keeping LMW antigens in solution. mAbs displayed dissociation constant (K_D_) values ranging more than 3 logs from 0.08 nM to 200 nM, which corresponds to low to very high-affinity antibodies (Fig. 3). We identified mAbs with a 1nM affinity or better for all ribotypes, except for RT014 that was only bound by mAb QE2 with a 9nM affinity. Noticeably, cross-specific mAbs displayed different affinities for their targets, with systematically one ribotype bound with at least a 10-fold better affinity, except for mAb VA10 that bound LMW-RT001 and LMW-RT027 with comparable affinities.

**Figure 3:**
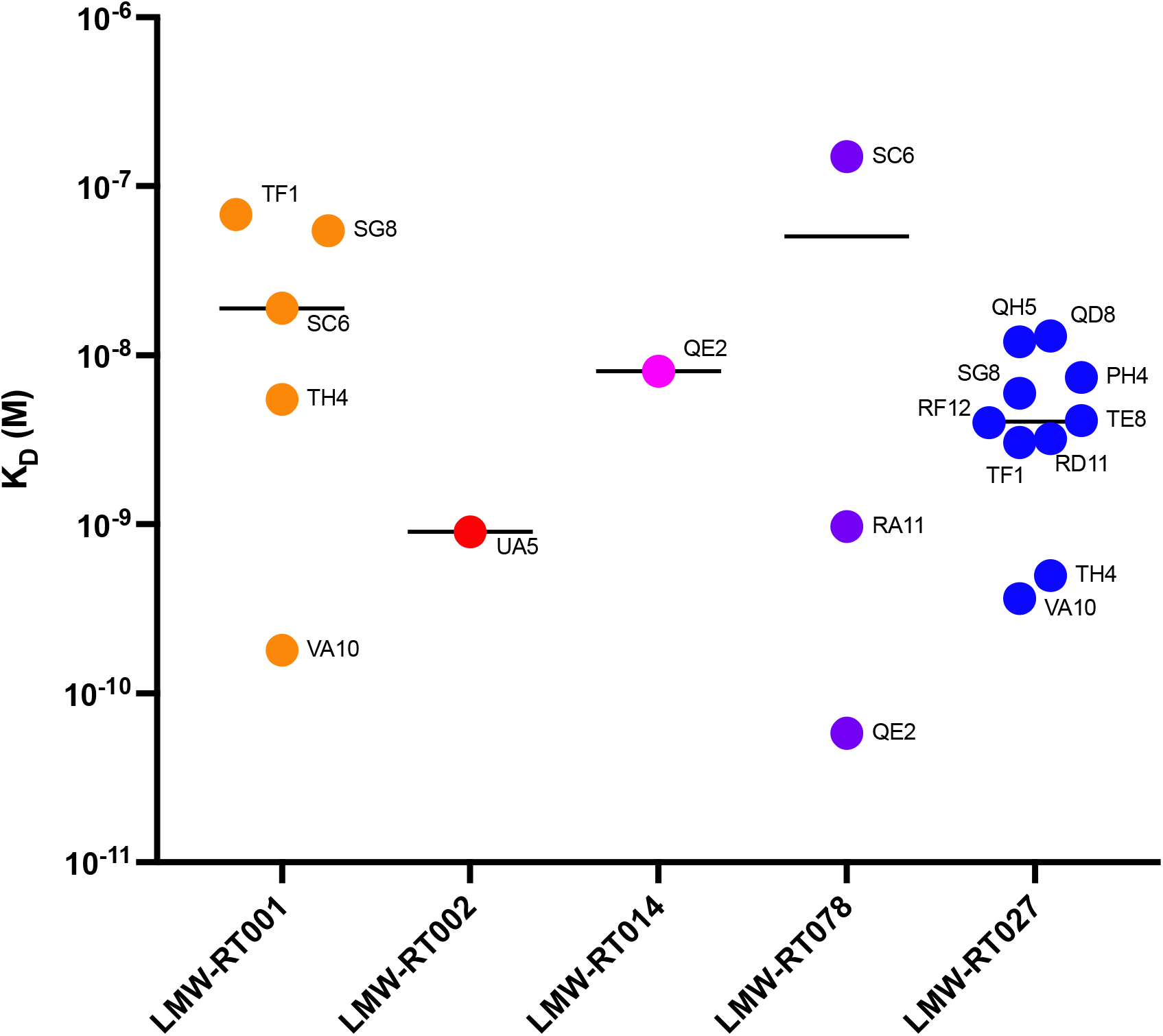
Affinities of mAbs for the LMW of five clinical ribotypes. Dissociation constant (K_D_) values measured by BLI. Each dot represents the K_D_ value of one mAb (mAb name indicated) interacting with one LMW among LMW-RT001, LMW-RT002, LMW-RT014, LMW-RT078 and LMW-RT027. Black bars represent median KD values of the group of mAbs binding one ribotype.

As SlpA is the main component of the *C. difficile* surface, we investigated if this series of mAbs could also bind LMW when exposed naturally at the bacterial surface. Fixed *C. difficile* from the different ribotypes were used for bacterial flow cytometry (Fig. 4a). Each ribotype could be significantly bound by at least one mAb. Consistent with the ELISA results (Fig. 2), monospecific anti-LMW mAbs, the LMW-RT027-specific mAbs (PH4, QD8, QH5, RD11 and TE8) and anti-LMW-RT078-specific mAbs (RA11), bound to *C. difficile* RT027 and RT078 whole bacteria, respectively. However, cross-specific mAbs bound a restricted number of ribotypes by bacterial flow cytometry (Fig. 4a) compared to ELISA (Fig. 2), indicating that their epitopes are hidden or inaccessible, or that their affinity is not sufficient for flow cytometry detection. Indeed, 3 out 8 cross-specific mAbs showed restricted binding profile using flow cytometry, *e*.*g*., QE2 mAb bound 4 distinct recombinant LMW ribotypes by ELISA but only 2 *C. difficile* ribotypes using flow cytometry. Table 1 summarizes the binding profiles of these mAbs to the LMW recombinant proteins and the LMW exposed at the bacterial surface for the five clinical ribotypes RT001, RT002, RT014, RT078, RT027.

**Table 1:** Summary table of mAbs binding profiles to LMW recombinant proteins and LMW expressed at the bacterial surface of *C. difficile* bacteria for five clinical ribotypes. E indicates binding by ELISA and F binding by flow cytometry. Blanks indicate absence of binding.

| Antibody | RT |  |  |  |  |
| --- | --- | --- | --- | --- | --- |
|  | 001 | 002 | 014 | 078 | 027 |
| PH4 | E |  |  | E/F | E/F |
| QD8 | E |  |  | E | E/F |
| QH5 | E |  |  |  | E/F |
| RD11 |  |  |  |  | E/F |
| RF12 | E |  |  | E | E/F |
| SG8 | E/F |  |  |  | E/F |
| TE8 | E |  |  |  | E/F |
| TF1 | E/F |  |  |  | E/F |
| TH4 | E/F |  |  |  | E/F |
| VA10 | E/F |  |  |  | E/F |
| QE2 | E |  | E/F | E/F | E |
| RA11 |  |  |  | E/F |  |
| SC6 | E/F |  |  | E/F |  |
| UA5 | E | E/F |  |  |  |

**Figure 4:**
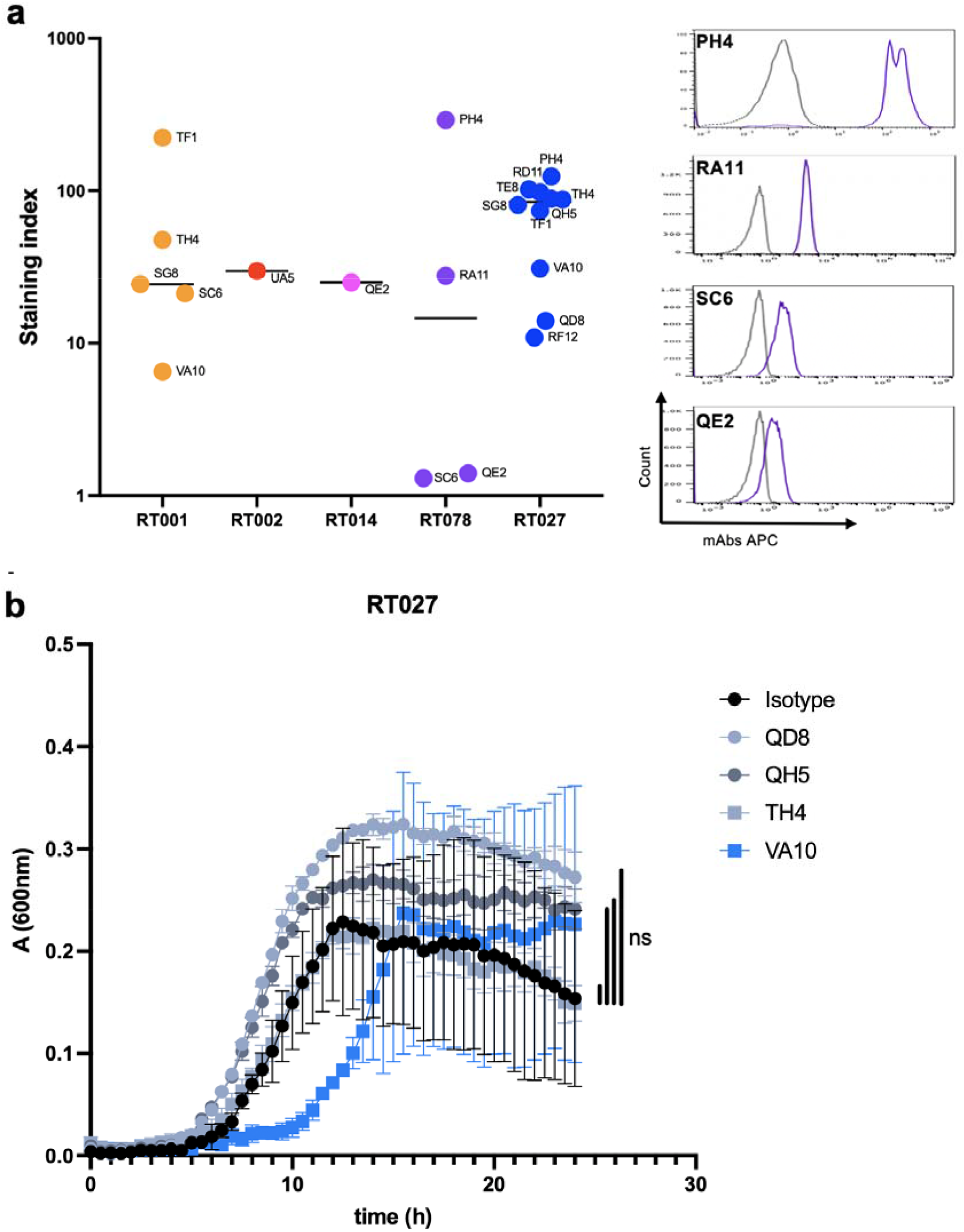
Binding of mAbs to LMWs expressed at the surface of *C. difficile* bacteria. (**a**) *Right*: Flow cytometry analysis of mAbs binding to LMW of indicated *C. difficile* ribotypes. Results are displayed as staining index (*refer to methods section*). *Left*: representative histograms for staining of strain RT078 by mAbs PH4, RA11, SC6 and QE2 are shown. (**b**) Growth of *C. difficile* strain RT027 in BHISG medium incubated with indicated anti-LMW027 mAb or with an unspecific IgG (isotype). Growth was followed continuously over 24h. Each dot represents the mean of three technical replicates, and the bars indicate standard deviations. ns: non-significant.

Finally, we studied the impact of LMW binding by the anti-LMW-RT027 mAbs in an *in vitro* growth assay on *C. difficile* strain 027. Two monospecific mAbs for LMW-RT027 (QD8 and QH5) and two cross-specific mAbs (VA10 and TH4) were tested for their impact on growth. Growth was followed over 24 hours with an isotype control IgG and showed an exponential phase followed by a plateau (Fig. 4b). Anti-LMW-RT027 did not significantly alter growth, even though mAb VA10 tended to delay growth, and mAb QD8 and, to a lesser extent, mAb QH5, tended to increase growth.

## DISCUSSION

Herein, we report the first monoclonal antibody collection that targets a surface protein of *C. difficile*. Due to sequence variability in the low-molecular weight subunit of surface layer protein A, this mAb collection allows the detection of 5 different ribotypes of clinical interest. More than half the mAbs bound selectively to the bacterial surface of one of these ribotypes, whereas the cross-reactive mAbs bound to two different ribotypes. The relatively high affinity of the interaction (nanomolar range) allows to envision using these mAbs for various assays such as ELISA, flow cytometry, microscopy, or histology assays.

In this study we chose to immunize mice with the low-molecular weight subunit of surface layer protein A as it represents a major antigen of the *C. difficile* surface^27^. Although we found by alignment stretches of conserved residues between the five ribotype sequences we used^18^, we could not identify any antibody cross-binding all five strains. The most cross-reactive anti-LMW mAbs recognized by bacterial flow cytometry only two different ribotypes. This suggests that conserved epitopes between LMW of different strains may not be dominant epitopes in terms of immunogenicity or may be hidden or poorly accessible to antibodies. Indeed, conserved amino acids have been implicated in the interaction between the LMW and the High Molecular Weight subunits which face inward toward the bacterial cell wall^28^ and are therefore probably inaccessible to antibodies.

Mice were immunized sequentially with five different LMWs and boosted with a mix of all of them, leading to identification of mAbs to each of them. Varying the order of different LMWs in the immunization scheme did not significantly alter antibody titers for the various LMWs, except for LMW-RT001 when injected with a farther ribotype. Antibodies binding SlpA have also been detected in the sera of patients infected with *C. difficile*, suggesting that, indeed, SlpA or its LMW subunit are immunogenic. Even though the knock-in mice we used produce antibodies with human variable domains^23,24^, thus potentially resembling those found in infected patients, we did not identify antibodies that significantly alter bacterial growth in our *in vitro* assays. It remains unclear whether such antibodies exist in patients in remission or if other mechanisms are at play. Interestingly, 30% of relapsing *C. difficile* infections are not due to the initial infecting strain but to a different strain, acquired from an exogenous source^29^. Whether the sequence variability of LMW among *C. difficile* ribotypes is involved in this recurrence and escape from the host immune response remains to be investigated.

This novel series of anti-*C. difficile* mAbs contains three anti-LMW mAbs specifically recognizing hypervirulent ribotypes RT027, bound by mAb TE8, RT078 bound by mAb RA11, and RT002 bound by mAb U5A. These three ribotypes have been associated with poor outcomes after infection^6,30,31^. Beyond *C. difficile* 630, the most studied *C. difficile* ribotype, this set of mAbs could help to study ribotypes RT027, RT078 and RT002 by resorting to various assays (ELISA, flow cytometry, microscopy, histology, blotting). One could even propose targeted treatments, by coupling antibiotics to these mAbs (aka Antibody-Drug Conjugates, ADC) to reduce antibiotic doses.

Our study however has limitations. While it has recently been reported, using whole-genome sequencing, that diversity exists within a given ribotype^32^, we only tested five ribotypes of *C. difficile*, each derived from a single clinical isolate. Therefore, more clinical isolates now remain to be tested to determine whether mAb specificity encompasses all known strains in each ribotype. Moreover, we only tested cross-specificity towards a limited panel of ribotypes. It remains to be deciphered if these mAbs cross-react with other *C. difficile* ribotypes not studied in this work.

To our knowledge, these mAbs represent the first collection of antibodies against *C. difficile* surface protein SlpA. These mAbs bind LMW from different clinically relevant strains *i*.*e*., LMW-RT001, LMW-RT002, LMW-RT014, LMW-RT027 and LMW-RT078. These mAbs represent interesting probes to better understand *C. difficile* infection, pathogenesis, and epidemiology.

## MATERIALS AND METHODS

### Mice

Knock-in mice expressing human antibody variable genes for the heavy (V_H_) and light chain (V_L_) (VelocImmune) were described previously^23,24^ and provided by Regeneron Pharmaceuticals to be bred at Institut Pasteur. All animal care and experimentation were conducted in compliance with the guidelines. The study, registered under #210111 was approved by the Animal Ethics committee CETEA (Institut Pasteur, Paris, France) and by the French Ministry of Research.

### Production of recombinant LMW proteins

Recombinant *C. difficile* LMW-SLPs (LMW-RT001, LMW-RT002, LMW-RT014, LMW-RT078, LMW-RT027, LMW630^25^) were produced as N-terminal 6xHis-tagged proteins from plasmid pET-28a(+) (TwistBiosciences, #69864). Plasmids were transformed into *E. coli* strain DE3 and grown in NZY auto-induction lysogeny broth (LB) medium (NZYtech, #MB180). Bacteria were harvested by centrifugation and lysed using Cell Disruptor (Constant System) at 1.3 kbar. Recombinant LMW-SLP proteins from the soluble fraction were purified by affinity chromatography on Histrap FF crude 1mL columns (Cytiva life science, #29048631) followed by size exclusion chromatography on HiLoad 16/600 Superdex 75 pg (Cytiva life science, #28989333) using an AKTA pure (Cytiva life science). All proteins were stored in 50 mM sodium phosphate buffer pH 7.8, 300mM NaCl prior to analysis or long-term storage.

### Production of LMW-specific monoclonal antibodies

VelocImmune mice were injected i.p. at day 0, 21 and 42 with 50 μg of each of five recombinant LMWs in alum mixed with 200 ng/mouse pertussis toxin (Sigma-Aldrich, #70323-44-3). ELISA was performed to measure serum responses to antigen (see methods below) and the 3 best immunized animals were boosted with the same antigen mix. Four days later, splenocytes were fused with myeloma cells P3X63Ag8 (ATCC, #TIB-9) using ClonaCell-HY Hybridoma Kit according to manufacturer’s instructions (StemCell Technologies, #03800). Culture supernatants were screened using ELISA (see below) and antigen-reactive clones were expanded in RPMI-1640 complemented with 10% IgG-free Fetal Calf Serum (Sigma-Aldrich, #F1283) into roller bottles (Sigma-Aldrich, #CLS431344) at 37°C. After 14 days, supernatants were harvested by centrifugation at 2500 rpm for 30 min and filtered (0.2 μm). Antibodies were purified by protein A affinity chromatography (AKTA pure) as described previously^33^.

### ELISA assays

Maxisorp microtiter plates (Dutscher, #055260) were coated with a total of 0.3 μg per well of LMW recombinant proteins in carbonate-bicarbonate buffer (pH 9.6) for 2 hours at room temperature (RT). Free sites were blocked by a 2-hour incubation at RT with PBS 1% BSA. Plates were washed three times with PBS 0.05% Tween 20 (PBS-T) before being coincubated with serum, supernatants, or monoclonal antibodies at different concentrations (from 10^−6^ μg/mL to 10 μg/mL) for 1h at RT. After five washes, goat anti-mouse IgG-Fc fragment HRP conjugated antibody (Bethyl, dilution 1:20,000, #A90-131P) was added for 1h at RT followed by incubation with OPD (o-phenylenediamine dihydrochloride) revelation substrate for 10 min (Sigma-Aldrich, #P8287). Absorbances were analyzed at 492 vs 620 nm on an ELISA plate reader (Berthold).

### Bio-layer interferometry

Biolayer interferometry assays were performed using Anti-Mouse Fc Capture biosensors on an Octet Red384 instrument (ForteBio, #18-5088). Monoclonal antibodies (10 μg/mL) were captured on the sensors at 25°C for 1,800 seconds. Biosensors were equilibrated for 10 minutes in 1x-PBS, 0,05% Tween 20, 0.1% BSA (PBS-BT) prior to measurement. Association was monitored for 1,200s in PBS-BT with LMW at a range of concentrations from 0.01 nM to 500 nM followed by dissociation for 1,200s in PBS-BT. Traces were reference sensor (sensors loaded with an unspecific mAb) subtracted and curve fitting was performed using a global 1:1 binding model in the HT Data analysis software 11.1 (ForteBio), allowing to determine K_D_ values.

### Flow cytometry assays

mAb binding to whole bacteria was assessed by bacterial flow cytometry, as previously described^34^. Briefly, fixed *C. difficile* (10^6^/condition) were stained with 5 μM Syto9 (Thermo Fisher Scientific, #S34854) in 0.9% NaCl for 30 min at RT. Bacteria were washed (10 min, 4,000g, 4°C) and resuspended in 1X PBS, 2% BSA and 0.02% Sodium Azide (PBA). Monoclonal antibodies were pre-diluted in PBA at 20 μg/mL and incubated for 30 min at 4°C. Bacteria were washed, and AF647 AffiniPure goat anti-mouse IgG (H+L) antibody or isotype control (Jackson ImmunoResearch, #115-605-003) were incubated for 30 min at 4°C. After washing, bacteria were resuspended in sterile 1X-PBS. Flow cytometry acquisition was performed on a MacsQuant cytometer (Miltenyi) and analyzed on FlowJo software (BD Biosciences). Staining index was calculated by subtracting the MFI of the isotype from the MFI of each condition with the anti-LMW mAbs, then divided by the MFI of the isotype.

### Bacterial strains and culture conditions

Clinical isolates of *C. difficile* RT001, RT002, RT014, RT027, RT078 were provided by The French National Reference Laboratory for *C. difficile*. Strains were grown anaerobically (5% H2, 5% CO2, 90% N2) in TY medium (30 g/L tryptone, 20 g/L yeast extract). Bacteria were fixed in 4% paraformaldehyde (PFA) for 30 min and resuspended in 1X PBS, 10% glycerol before being stored at -80°C. All media and chemicals were purchased from Sigma-Aldrich.

### Growth assays

Overnight *C. difficile* cultures were grown in TY broth and sub-cultured to an Optical Density at 600 nm (OD600nm) of 0.05 in 200 μL of BHISG in 96-well flat bottom plates (Merck, #Z707902). Bacterial growth was followed for 24h or 18h with OD600nm measurements every 30 min using GloMax Plate Reader (Promega). Anaerobia was maintained with a O_2_ less sealing film (Sigma-Aldrich, #Z380059).

### Statistical analysis

Growth and ELISA assays values were analyzed in Prism 8.0 (GraphPad, San Diego, CA). Statistical analysis was performed using two-way ANOVA test. A p value ≤0.05 was considered significant.

## ACKNOWLEDGEMENTS

We thank Stéphane Pêtres (Production and Purification of Recombinant Proteins Facility, Institut Pasteur, 75015 Paris, France) for his advice on protein purifications. This work was funded by Fondation Janssen Horizon, the Institut National de la Santé et de la Recherche Médicale (INSERM) and the Institut Pasteur. LH is a doctoral fellow of Sorbonne Université. DS was a recipient of a poste d’accueil 2017 Institut Pasteur – Assistance Publique des Hôpitaux de Paris (APHP). Work in the G. Gorochov’s team is supported by Institut National de la Santé et de la Recherche Médicale (INSERM), Sorbonne Université, Fondation pour la Recherche Médicale (FRM), Paris, France, program ‘‘Investissement d’Avenir’’ launched by the French Government and implemented by the Agence Nationale de la Recherche (ANR) with the reference COFIFERON ANR-21-RHUS-08, by EU Horizon HLTH-2021-DISEASE-04 UNDINE project, by programme DIM Ile de France thérapie cellulaire et génique, by Fondation pour la Recherche Médicale, Paris, France (Programme Equipe FRM 2022) and by the Département Médico-Universitaire de Biologie et Génomique Médicales (DMU BioGen), APHP, Paris, France.

## AUTHORSHIP CONTRIBUTIONS

Experimental design, LH, DS and PB; Conducting experiments, LH, BI, OG; Data analyses and discussions: LH, PE, FB, BD, LM, GG, DS and PB. Writing (original draft), LH, DS and PB; Writing (review and editing), all authors.

## COMPETING INTERESTS

Unrelated to the submitted work, P.B. received consulting fees from Regeneron Pharmaceuticals. The other authors declare no competing interests.

## FIGURES

**Figure S1.**
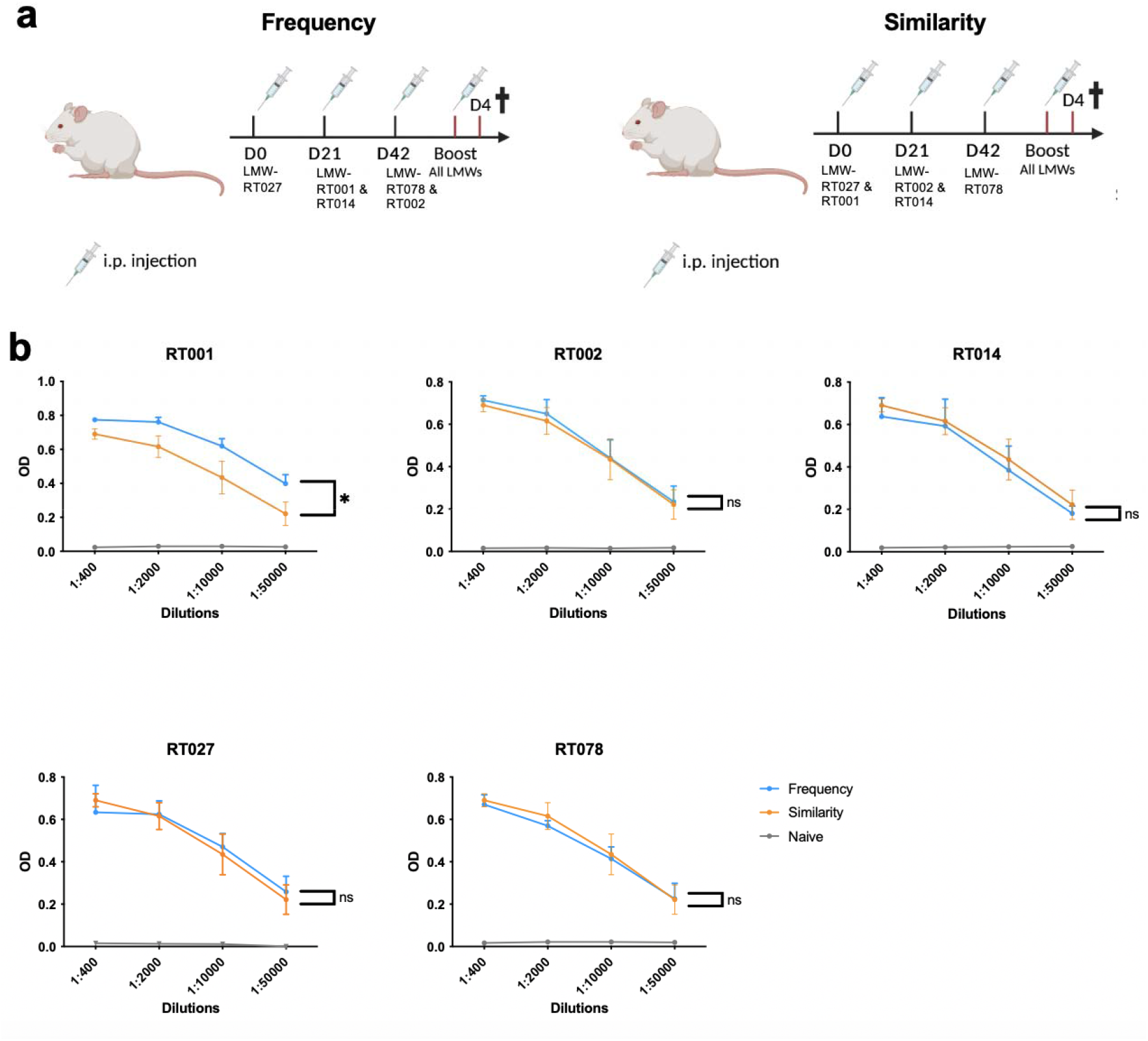
Comparison of two immunization protocols using recombinant LMWs. Mice were immunized following two different protocols termed “similarity” and “frequency”. (**a**) In the “Frequency” protocol, mice are immunized with LMWs in the order of their frequency in current CDI, and boosted with a mix of all five LMWs. In the “Similarity” protocol, mice are immunized with two highly similar LMW the same day, and boosted with a mix of all five LMWs. (**b**) Dose response of sera titers of immunized mice from the protocols depicted in (a) are measured by ELISA against the indicated LMW ribotype. Data are presented as mean values (±SD) for each group of mice (n=5). ns: non-significant; *: p<0.05. Black curves represent sera from naive mice prior immunization.

